# Neutools: A collection of bioinformatics web tools for neurogenomic analysis

**DOI:** 10.1101/429639

**Authors:** Fan Gao, Hugh P. Cam, Li-Huei Tsai

## Abstract

With a large and growing number of neurogenomic data, from epigenetic, transcriptomic to proteomic data deposited to the public domains, visualization and mining of these data alongside one own data have become extremely useful for identifying potential genetic targets and/or biological pathways to further validate and characterize in model systems. Here, a series of easy-to-use web tools (**Neutools**) were developed using Shiny/R codes for neuroscientists to perform basic bioinformatics data analysis and visualization. Specifically, **NeuVenn** calculates and plots overlap statistics for multiple input gene sets; **NeuGene** and **NeuChIP** visualize gene expression data and histone ChIP-Seq data generated from brain related tissue and cell culture, respectively. **NeuVar** annotates human brain GWAS variants and epigenetic features based on user-specified genes and regions. **Neutools** are freely available resources for all academic users to use.

## Introduction

Studies of biological systems increasingly rely on advances in high-throughput omics approaches. To a certain degree, molecular biology is gradually becoming a “big-data science.” Gigabytes to terabytes of raw data related to genomics, epigenetics, transcriptomics, proteomics, and metabolomics are routinely generated in studies and deposited to public repositories. Analysis of published omic datasets generated from the same or similar model systems could rapidly yield insights leading to testable hypothesis not possible with traditional literature search or limited empirical work. Thus, the development of new and freely available bioinformatics tools and resources becomes essential to support big-data analysis.

Large-scale population genetics studies have identified numerous risk genes/SNPs associated with different neurological disorders. Region and cell-type specific isolation of brain tissue and neuronal cell culture together with high-throughput profiling techniques in genomics, epigenomics, transcriptomics, and proteomics have been well-received approaches for systematic studies. However, the high-throughput data released from individual studies are typically processed with different bioinformatics pipelines and difficult to visualize (stored either as raw data or processed table formats). Therefore, new user-friendly bioinformatics tools with easy-touse query interface, graphic visualization of queried data, and basic statistics tables of the query would be welcome addition for most neuroscientists not familiar with bioinformatics languages.

In the current study, a collection of Shiny/R based web tools named **Neutools** have been developed for neuroscientists to 1) analyze gene overlap statistics (**NeuVenn**); 2) visualize gene expression data generated from high-throughput transcriptomics or proteomics (**NeuGene**); 3) display epigenetic histone ChIP-Seq data around gene promoters (**NeuChIP**); 4) annotate human brain GWAS variants and epigenetic features on specified regions (**NeuVar**). By querying against the curated lists or the processed omic data, these tools provide basic bioinformatics functionality for discovering various molecular signatures in the nervous system.

## Material and methods

Human and mouse next-generation sequencing raw data were collected mostly from public data repositories such as Gene Expression Omnibus (GEO, https://www.ncbi.nlm.nih.gov/geo/), Roadmap Epigenetics Mapping Consortium (http://www.roadmapepigenetics.org), ENCODE Portal (https://www.encodeproject.org), and AMP-AD Knowledge Portal (https://www.synapse.org/#!Synapse:syn2580853). Transcriptomic RNA-Seq data were aligned to and quantified using either GENCODE-v24 human annotation or GENCODE-vM9 mouse annotation (https://www.gencodegenes.org). Gene expression FPKM matrices were generated from the default Cufflinks pipeline^1^ and saved for query purpose. ChIP-Seq data were aligned to either human genome assembly (hg19) or mouse assembly (mm9) using BWA aligner^2^.

Duplicate reads were removed by Samtools^3^. Genome-wide signal intensity traces were generated in bedgraph format, and processed using a custom PERL script to calculate signal intensity matrices around the transcription start sites of the reference genes for query purpose. Other processed data were collected from individual data portals (see details in RESULTS).

The web interfaces for data query were written in Shiny/R codes. Shiny is an easy-to-use open source R package that builds elegant interactive web apps directly from R codes. Neutools were designed with two display panels, with the left panel for query and the right panel for output (**Figure 1**). Users can check software update by clicking the triangle at the top right corner of the app page (**Figure 1**). A short introductory page with a start-up guide is also provided for each software tool.

**Figure 1.**
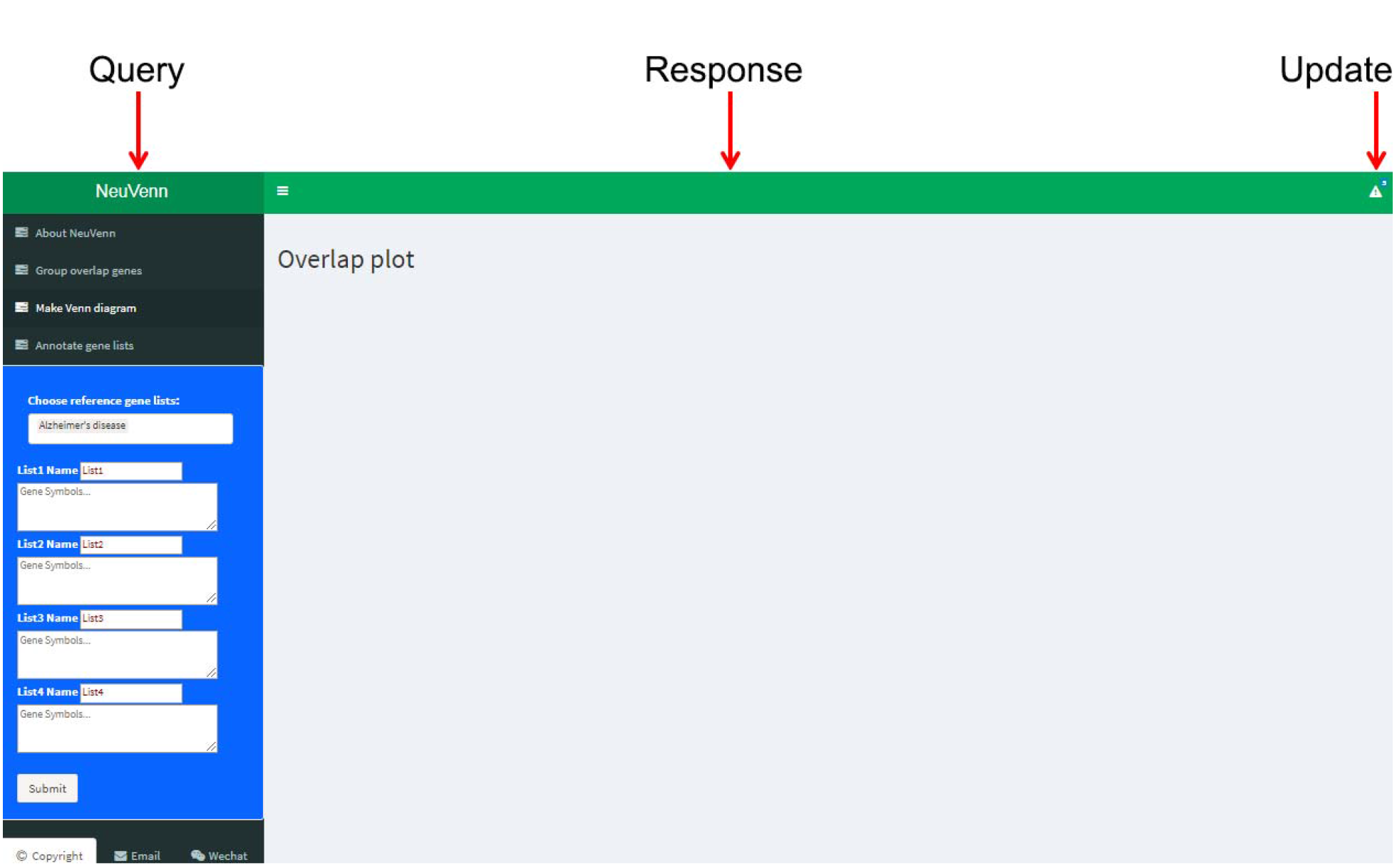
Shiny/R web design for Neutools. Neutools use the Bootstrap theme embedded in Shiny. The input (query) and output (response) sections are on the same web page, enablingthe users to easily modify the search terms and check the interactive results.

The benchmark tests of the developed web tools on a DELL POWEREDGE 2950 server showed reasonable response time in seconds for the majority of functionalities, and in one or two minutes for the rest of the applications.

## Results

**NeuVenn** (http://bioinfo5pilm46.mit.edu:318/neuvenn/) calculates and plots overlap statistics for several user defined gene sets (human and mouse gene symbols as input). This software generates overlap gene lists and statistical p-values (Fisher’s exact test) (**Figure 2 B**), and makes Venn diagrams for overlap gene counts (**Figure 2 A, C**). In addition, it annotates user gene sets by checking overlap statistics with the curated brain disease/disorder and cell-type specific gene sets^4–10^ (**Figure 2 D**). NeuVenn is useful in determining whether a group of differentially expressed genes (DEGs) from RNA-seq experiments seen under one condition (e.g., genetic or chemical perturbation) is statistically overlapped with another group of DEGs under a different condition. NeuVenn can simultaneously compare up to four different groups of gene-sets.

**Figure 2.**
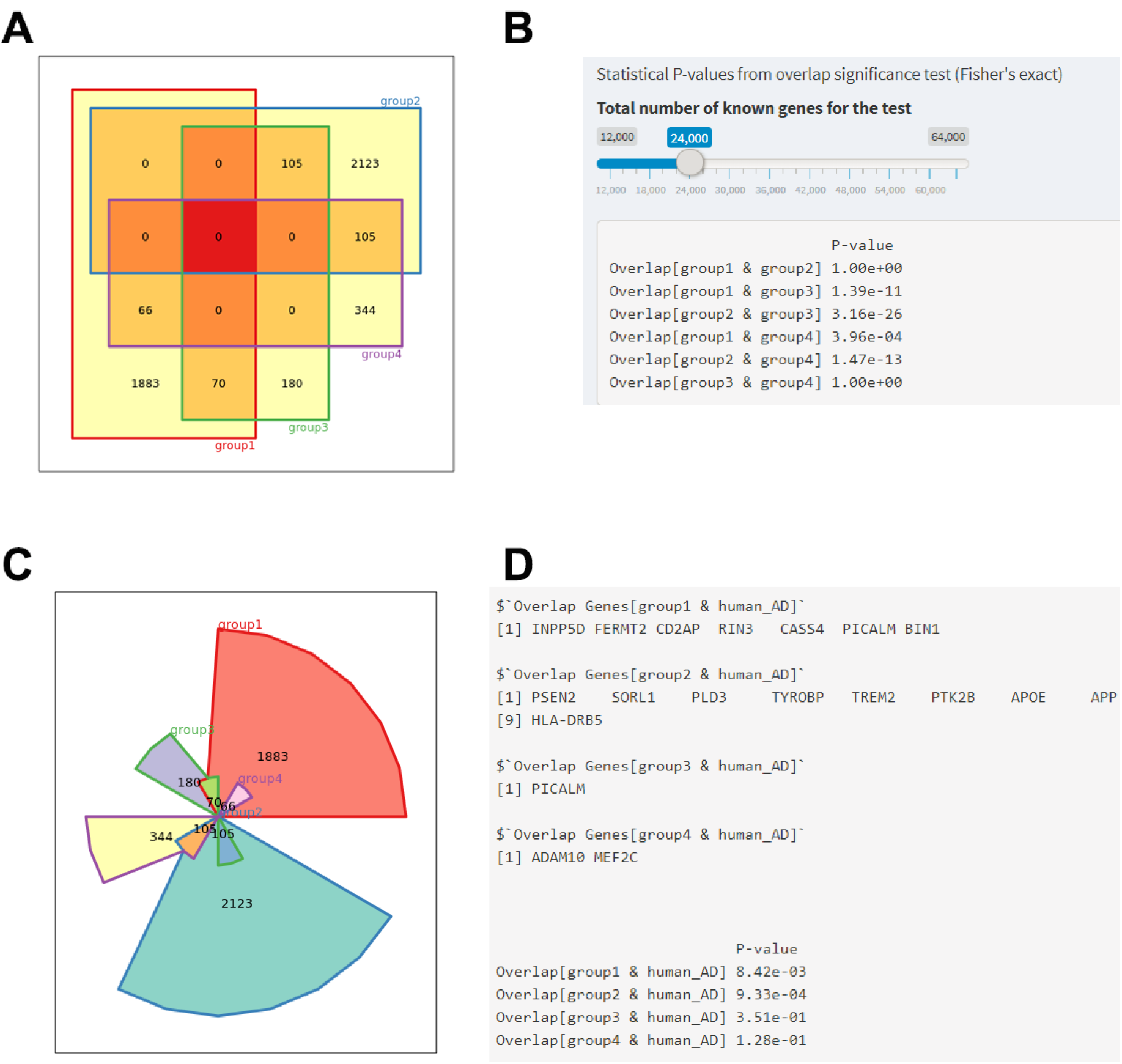
NeuVenn demo results. Four different gene groups were uploaded to generate unweighted (**A**) and weighted (**C**) Venn Diagrams. Overlap statistical p-values were generated using Fisher’s exact test (**B**). Queried genes that overlap with the curated brain disease/disorder and cell-type specific genes were identified (**D**).

**NeuGene** (http://bioinfo5pilm46.mit.edu:318/neugene/) visualizes gene expression data generated from brain tissue (region specific or cell-type specific) and in-vitro cell culture. This software searches a specific category of expression data (such as Alzheimer’s disease, Huntington’s disease, brain aging process, different chemical perturbations, neuronal activity and others) in the database and generates bar charts to compare gene expression pattern across different sample groups (**Figure 3**). Transcriptomic data such as those from RNA-Seq are calculated as FPKM values, and high-throughput proteomic data are quantified as log_2_[LFQ] values. In addition, **NeuGene** provides an option to annotate gene types for your query genes. NeuGene is highly useful for a quick look-up of RNA or protein expression for a small set of genes (up to ten queries) of interests in specific brain region and cell-type under certain conditions (aging, neurodegenerative diseases, genetic or chemical perturbations).

**Figure 3:**
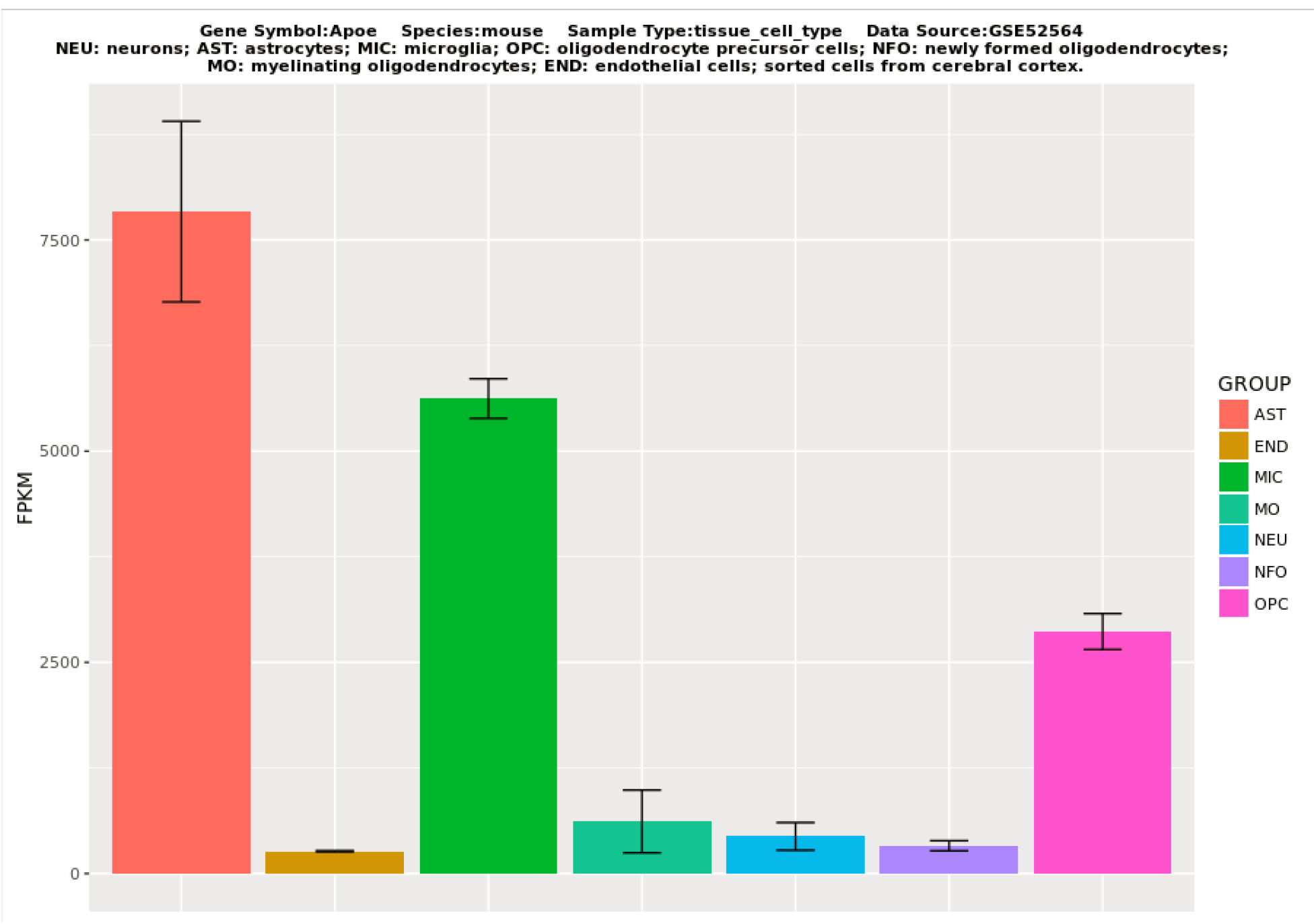
NeuGene demo result. Gene expression levels of mouse ApoE across different brain cell types were visualized in the bar chart. Group mean FPKM values and the stand errors were shown in the plot.

**NeuChIP** (http://bioinfo5pilm46.mit.edu:318/neuchip/) provides graphical presentation of human and mouse ChIP-Seq data of histone modifications generated from brain tissue and invitro cell culture. This software searches specific histone mark(s) in the database and generates signal intensity aggregation plots and heatmaps (**Figure 4 A, B**). Histone signals around transcription start sites of the query gene list(s) are visualized for cross-group comparison. In addition, **NeuChIP** provides 1) chromosomal location and transcriptional orientation of your gene list(s); 2) density ideogram of your gene list(s) on the chromosomes (**Figure 4 C**). Similar to NeuGene, NeuChIP is adept at providing quick look-ups of epigenomic information for a particular gene promoter of interest with scalable resolution in specific brain region and cell-types under certain conditions (e.g., neurodevelopment, neurodegenerative diseases).

**Figure 4:**
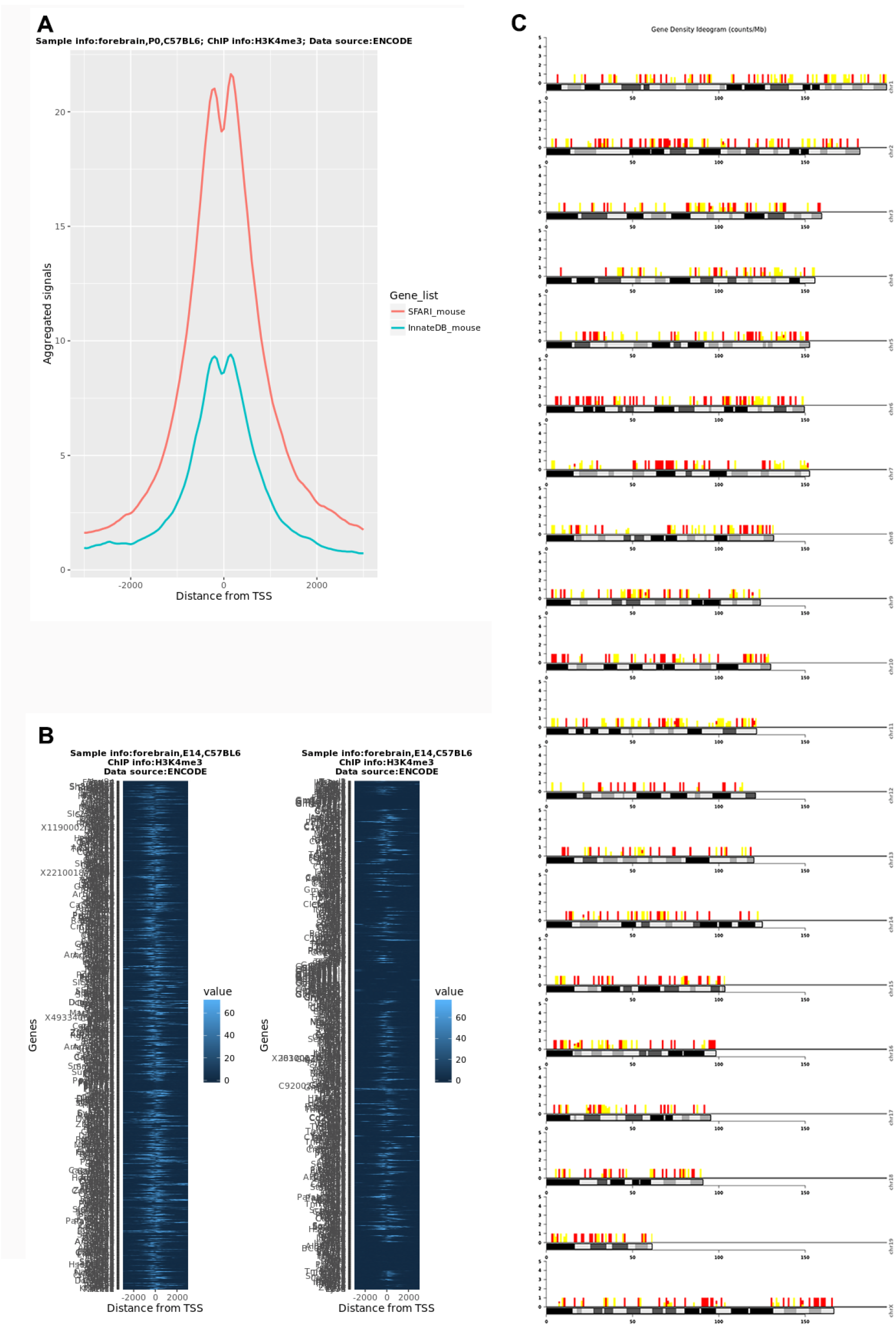
NeuChIP demo results. Two mouse gene sets (SFARI ASD-related genes and InnateDB immune-related genes) were uploaded to generate histone H3K4me3 ChIP-Seq aggregated intensity map (**A**) and heatmap (**B**). Gene density of the queries on different chromosomes was visualized in density ideogram (**C**).

**Figure 5:**
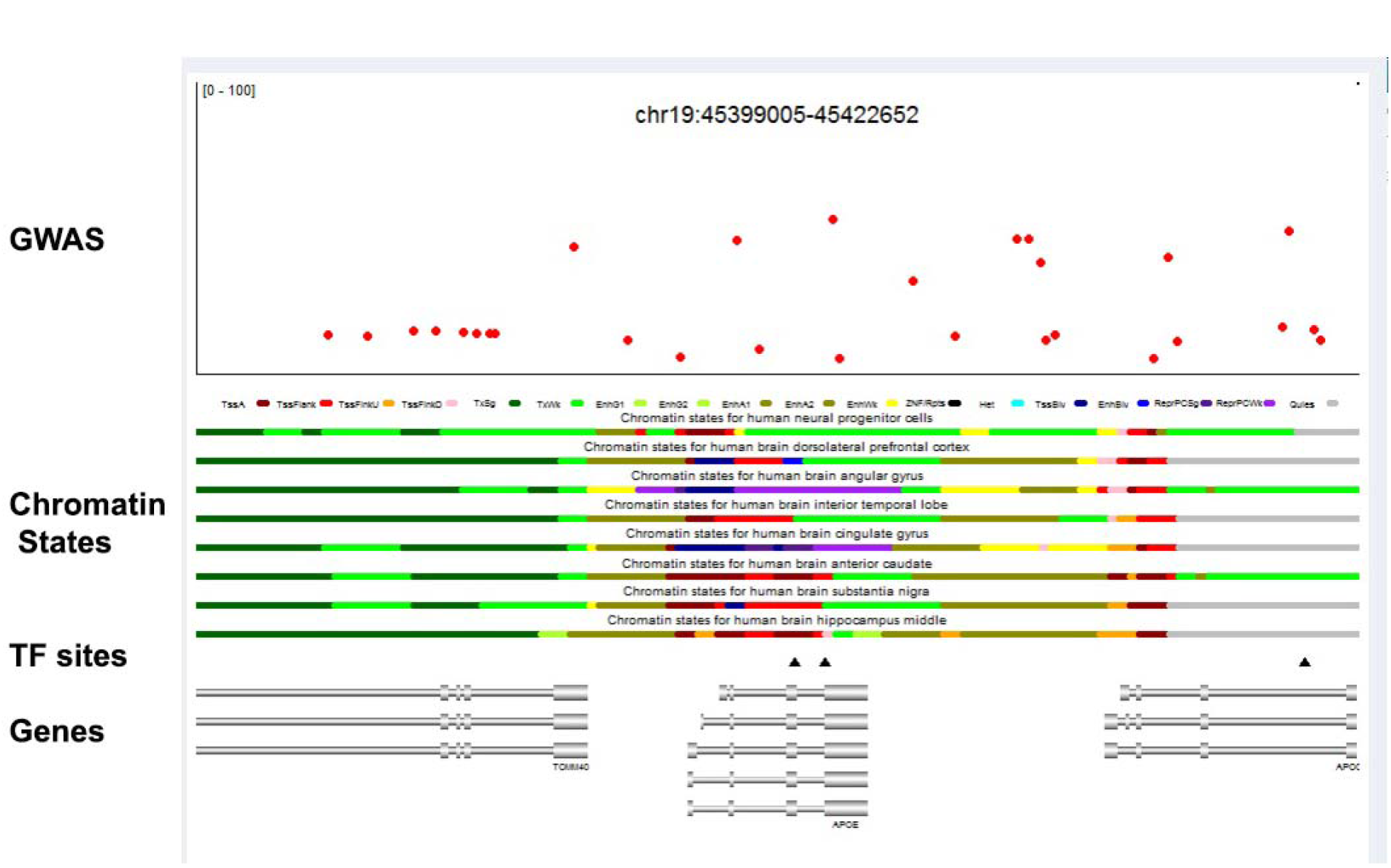
NeuVar demo result. Genomic regions surrounding *APOE* gene locus were annotated with genetic neural GWAS hits (top panel), epigenetic brain chromatin states and conserved transcription factor binding sites (middle panel), and gene structures (bottom panel).

**NeuVar** (http://bioinfo5pilm46.mit.edu:318/neuvar/) annotates human brain GWAS variants and epigenetic features of user-specified genes and genomic regions (up to ten queries). This software generates interactive tables with sort, search, filter functions for the GWAS hits and epigenomic features that overlap with your queried human gene and genomic regions.

**NeuVar** database includes the following datasets:

1. Human brain GWAS data collected from different data repositories, including International Genomics of Alzheimer’s Project (IGAP), Alzheimer’s Disease Genetics Consortium (ADGC), Social Science Genetic Association Consortium (SSGAC), Psychiatric Genomics Consortium (PGC), and other studies.
2. NIH ROADMAP chromatin states defined by the combination of six histone marks (H3K4m3, H3K4me1, H3K36me3, H3K27me3, H3K9me3 & H3K27ac) in different human brain regions (hippocampus middle, substantia nigra, anterior caudate, cingulate gyrus, interior temporal lobe, angular gyrus & dorsolateral prefrontal cortex) and human ES derived neural progenitor cells^11^.
3. NIH ENCODE conserved transcription factor binding sites (hg19_tfbsConsSites).

In addition, the software provides an option to upload user tracks (bed format, up to three) for the annotation plot.

With an interactive web design, selections/unselections of the contents on the result tables are displayed on the output canvas. Compared to **NeuChIP**, **NeuVar** is relatively slower but in return, it provides an integrated view of the rich landscapes of genetic information alongside several (at least eight) epigenomic tracks. For users interested in the genetic and epigenetic basis of neurodegenerative diseases and psychiatric disorders, the ability to visualize GWAS variants alongside epigenomic signatures would be highly useful for forming readily testable hypothesis regarding possible causal link between a GWAS variant and the corresponding epigenetic alteration and their roles in the disease or disorder.

In summary, a series of bioinformatics tools have been developed to meet the “big-data science” demand from molecular neuroscientists. With the powerful statistical tools embedded in R software, Shiny/R web development language has proven highly efficient for designing web-based scientific application tools for omics analysis of neurobiological research.

## Acknowledgements

This work was supported by JPB Foundation.

## Author Contributions

F.G. designed the work, developed the software and drafted the manuscript. H.P.C. and L.-H.T. reviewed and edited the manuscript. L.-H.T. conceived and supervised the work.

## References

1 Trapnell, C. et al. Differential analysis of gene regulation at transcript resolution with RNA-seq. Nat Biotechnol 31, 46–53, doi:10.1038/nbt.2450 (2013).

2 Li, H. & Durbin, R. Fast and accurate short read alignment with Burrows-Wheeler transform. Bioinformatics 25, 1754–1760, doi:10.1093/bioinformatics/btp324 (2009).

3 Li, H. et al. The Sequence Alignment/Map format and SAMtools. Bioinformatics 25, 2078–2079, doi:10.1093/bioinformatics/btp352 (2009).

4 Karch, C. M. & Goate, A. M. Alzheimer’s disease risk genes and mechanisms of disease pathogenesis. Biol Psychiatry 77, 43–51, doi:10.1016/j.biopsych.2014.05.006 (2015).

5 Schizophrenia Working Group of the Psychiatric Genomics, C. Biological insights from 108 schizophrenia-associated genetic loci. Nature 511, 421–427, doi:10.1038/nature13595 (2014).

6 Darmanis, S. et al. A survey of human brain transcriptome diversity at the single cell level. Proc Natl Acad Sci U S A 112, 7285–7290, doi:10.1073/pnas.1507125112 (2015).

7 Zhang, Y. et al. An RNA-sequencing transcriptome and splicing database of glia, neurons, and vascular cells of the cerebral cortex. J Neurosci 34, 11929–11947, doi:10.1523/JNEUROSCI.1860-14.2014 (2014).

8 Lee, C. Y. D. et al. Elevated TREM2 Gene Dosage Reprograms Microglia Responsivity and Ameliorates Pathological Phenotypes in Alzheimer’s Disease Models. Neuron 97, 1032–1048 e1035, doi:10.1016/j.neuron.2018.02.002 (2018).

9 Seyfried, N. T. et al. A Multi-network Approach Identifies Protein-Specific Co-expression in Asymptomatic and Symptomatic Alzheimer’s Disease. Cell Syst 4, 60–72 e64, doi:10.1016/j.cels.2016.11.006 (2017).

10 Sharma, K. et al. Cell type-and brain region-resolved mouse brain proteome. Nat Neurosci 18, 1819–1831, doi:10.1038/nn.4160 (2015).

11 Roadmap Epigenomics, C. et al. Integrative analysis of 111 reference human epigenomes. Nature 518, 317–330, doi:10.1038/nature14248 (2015).

